# Vti1a/b support distinct aspects of TGN and *cis*-/medial Golgi organization

**DOI:** 10.1101/2022.06.26.497636

**Authors:** Danique M van Bommel, Ruud F Toonen, Matthijs Verhage

## Abstract

Retrograde trafficking towards the *trans*-Golgi network (TGN) is important for dense core vesicle (DCV) biogenesis. Here, we used Vti1a/b deficient neurons to study the impact of disturbed retrograde trafficking on Golgi organization and cargo sorting. In Vti1a/b deficient neurons, staining intensity of *cis*-/medial Golgi proteins (e.g., GM130 and giantin) was increased, while intensity of TGN-resident proteins (e.g., TGN38 and TMEM87A) was decreased. Levels and localization of DCV cargo markers and LAMP1 were altered. This phenotype was not caused by reduced Golgi membrane availability or absence of the TGN compartment. The phenotype was partially phenocopied by disturbing sphingolipid homeostasis, but was not rescued by overexpression of sphingomyelin synthases or the sphingolipid synthesis inhibitor myriocin. We conclude that Vti1a/b are important for distinct aspects of TGN and *cis*-/medial Golgi organization. Our data underline the importance of retrograde trafficking for Golgi organization, DCV cargo sorting and the distribution of proteins of the regulated secretory pathway.

## Introduction

The Golgi complex is the major sorting organelle in the secretory pathway. It receives newly synthesized proteins from the endoplasmic reticulum and recycled cargo from other organelles, such as endosomes, by retrograde trafficking. In neurons, cargo leaving the Golgi apparatus can be sorted into constitutive or regulated secretory vesicles. Dense core vesicles (DCVs) are the main carriers of the regulated secretory pathway and contain neuropeptides and neurotrophins that modulate synaptic activity and play a role in the regulation of diverse functions such as fear, memory and appetite (Bramham & Messaoudi, 2005; Comeras et al., 2019; Zi et al., 2008). Alterations in neuropeptide/neurotrophin levels have been associated with pathological conditions such as anxiety, posttraumatic stress disorder and depression (Alldredge, 2010; Sah & Geracioti, 2013). DCV cargo is synthesized at the endoplasmic reticulum and transported via the Golgi apparatus to the *trans-Golgi* network (TGN) (Kim et al., 2006). How DCV cargo is sorted into DCVs and away from other secretory pathways at the TGN is poorly understood. Recent studies emphasize the importance of retrograde trafficking from endosomes to the TGN for DCV biogenesis (Emperador-Melero et al., 2018; Ma et al., 2020), but the interplay between these pathways remains elusive.

Ma et al. show that DCV maturation in larval salivary glands of *Drosophila melanogaster* requires retrograde transport from early endosomes (Ma et al., 2020). In addition, Vti1a/b, SNARE proteins involved in endosome-to-TGN retrograde trafficking, are necessary for DCV cargo sorting and exit from the Golgi (Emperador-Melero et al., 2018). Neurons that are deficient in Vti1a/b contain fewer DCVs and staining intensity of DCV cargo is reduced. Further experiments show that DCV cargo accumulates in the Golgi and that the heterogeneous distribution of DCV cargo in the Golgi is lost. The Golgi apparatus is more compact, but its cisternae are more distended. While transport of DCV cargo from the endoplasmic reticulum to the Golgi apparatus is normal, export of regulated cargo and, to a lesser extent, constitutive cargo from the TGN is reduced. It was proposed that the retrograde endosome-to-TGN pathway is crucial for retrieval of proteins involved in DCV biogenesis to the TGN. Characterizing the Golgi defects caused by loss of Vti1a/b in neurons will provide insight into the requirements for DCV biogenesis.

Here, we studied the impact of disturbed retrograde trafficking on the organization of the Golgi apparatus and proteins involved in Golgi export. For this goal, we used primary hippocampal neurons of Vti1a/b deficient mice and analyzed the localization and levels of *cis-/* medial Golgi and TGN proteins, and DCV cargo using confocal microscopy. We confirmed that the Golgi of Vti1a/b deficient neurons is condensed and that DCV cargo accumulates in the Golgi. Staining of Golgi-localized proteins was abnormal and levels and localization of proteins associated with DCVs were disturbed. Staining intensity of LAMP1-positive organelles was also reduced and their shapes were altered. These defects were not caused by reduced Golgi membrane availability or the absence of a TGN compartment. The phenotype was partially mimicked by addition of ceramide or sphingosine to neurons, but not rescued by overexpression of sphingomyelin synthases or treatment with myriocin. We conclude that Vti1a/b are required for a normal *cis-/*medial Golgi and TGN and for a normal distribution of proteins involved in regulated secretion.

## Results

To study the impact of disturbed retrograde trafficking on Golgi organization and proteins involved in Golgi export, we used mouse hippocampal neurons from Vti1a/b double knock-out (DKO), with double heterozygous (DHZ) mice as a control (Emperador-Melero et al., 2018). First, to confirm and extend the previously observed Golgi abnormalities, we examined levels and localization of two *cis-/*medial Golgi markers, three *trans*-Golgi network (TGN) proteins, two DCV cargoes and two proteins implicated in DCV biogenesis.

### *cis*-/medial Golgi and TGN protein staining is altered in Vti1a/b DKO neurons

Previously, it was shown that the *cis*-Golgi marker GM130 stains a smaller area and its staining intensity is increased in the absence of Vti1a/b (Emperador-Melero et al., 2018). We confirmed these observations here (Fig. 1a-c). In addition, the area stained by giantin, a *cis*-/medial Golgi marker, was also smaller and its average staining intensity was higher in DKO neurons than in controls (Fig. 1d-f). To estimate the total amount of GM130 and giantin localized to the Golgi, we measured the integrated density of the markers by multiplying the mean gray value and the area of the region of interest. Even though the average staining intensities of GM130 and giantin were higher, the Golgi integrated densities of these proteins were lower (Sup. Fig. 1a,b). These data confirm previous data and suggest that the expression levels of *cis*-/medial Golgi marker proteins are lower in the Golgi of Vti1a/b DKO neurons.

**Fig. 1.**
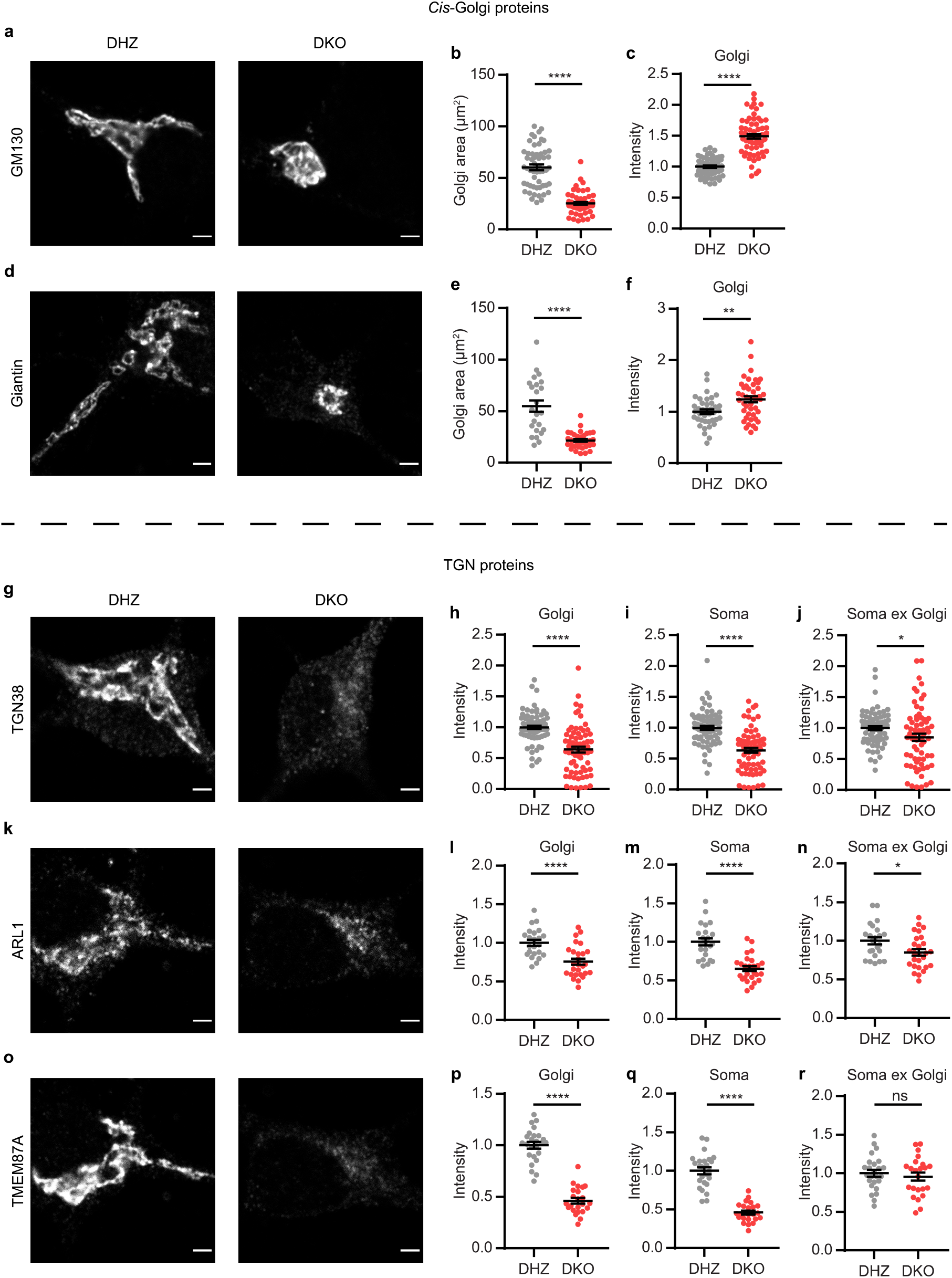
Staining intensity of *cis*-/medial Golgi proteins is increased in Vti1a/b deficient neurons, while TGN protein staining intensity is decreased. **a** Representative examples of neurons immunostained for GM130. **b** Golgi area based on GM130 staining is decreased in Vti1a/b DKO neurons compared to DHZ controls (DHZ: 58.7 ± 2.45, *n* = 64; DKO: 24.4 ± 1.26, *n* = 66; Mann Whitney test). **c** GM130 normalized staining intensity in the Golgi is increased (DHZ: 1 ± 0.0177, *n* = 65; DKO: 1.49 ± 0.039, *n* = 61; *t*-test). **d** Representative examples of neurons immunostained for giantin. **e** Golgi area based on giantin staining is smaller in Vti1a/b DKO neurons compared to DHZ neurons (DHZ: 54.6 ± 4.59, *n* = 29; DKO: 21.1 ± 1.25, *n* = 39; *t*-test). **f** Giantin normalized staining intensity in the Golgi is increased (DHZ: 1 ± 0.0445, *n* = 39; DKO: 1.24 ± 0.0572, *n* = 44; *t*-test). **g** Representative examples of neurons immunostained for TGN38. **h-j** TGN38 normalized staining intensity is decreased in the Golgi (DHZ: 1 ± 0.0294, *n* = 73; DKO: 0.641 ± 0.0476, *n* = 69; Mann Whitney test) (**h**), soma (DHZ: 1 ± 0.0322, *n* = 73; DKO: 0.636 ± 0.0416, *n* = 69; Mann Whitney test) (**i**), and soma excluding Golgi (DHZ: 1 ± 0.0324, *n* = 73; DKO: 0.851 ± 0.0581, *n* = 69; Mann Whitney test) (**j**). **k** Representative examples of neurons immunostained for ARL1. **l-n** ARL1 normalized staining intensity is decreased in the Golgi (DHZ: 1 ± 0.0387, *n* = 23; DKO: 0.756 ± 0.0397, *n* = 27; *t*-test) (**l**), soma (DHZ: 1 ± 0.0468, *n* = 23; DKO: 0.653 ± 0.0319, *n* = 27; *t*-test) (**m**), and soma excluding Golgi (DHZ: 1 ± 0.0464, *n* = 23; DKO: 0.851 ± 0.0422, *n* = 27; *t*-test) (**n**). **o** Representative examples of neurons immunostained for TMEM87A. **p-r** TMEM87A normalized staining intensity is decreased in the Golgi (DHZ: 1 ± 0.0323, *n* = 25; DKO: 0.462 ± 0.027, *n* = 23; *t*-test) (**p**) and soma (DHZ: 1 ± 0.0441, *n* = 25; DKO: 0.461 ± 0.0259, *n* = 23; *t*-test) (**q**), but not significantly different in the soma excluding Golgi (DHZ: 1 ± 0.0446, *n* = 25; DKO: 0.955 ± 0.053, *n* = 23; *t*-test) (**r**). Bars show mean ± SEM. *p < 0.05; **p < 0.01; ***p < 0.001; ****p < 0.0001. Scale bar is 2μm.

To examine the TGN, we used antibodies against TGN marker TGN38 and two proteins that localize to the TGN: TMEM87A and ARL1 (Fig. 1g,k,o). In contrast to *cis*-/medial Golgi markers, the average staining intensity of TGN proteins was decreased in the Golgi (Fig. 1h,l,p). The effect was largest for TMEM87A, for which the average staining intensity in the Golgi was reduced by ~50%, compared to control neurons. Similar to *cis*-/medial Golgi proteins, Golgi integrated density of all three TGN proteins was decreased in Vti1a/b DKO neurons (Sup. Fig 1c-e). This suggests that loss of Vti1a/b not only affects the level of *cis*-/medial Golgi proteins, but also of TGN proteins.

TGN38 shuttles between the plasma membrane and the TGN (McNamara et al., 2004; Reaves et al., 1993). Since Vti1a/b are required for retrograde trafficking to the Golgi, these proteins might need Vti1a/b to return to the Golgi. Loss of Vti1a/b could thus have resulted in altered TGN38 localization. As shown in figure 1i,m,q, the average staining intensity was lower in the whole soma of DKO neurons when compared to control neurons. In addition, the staining intensity in the soma excluding the Golgi was not different for TMEM87A and slightly decreased for TGN38 and ARL1 (Fig. 1j,n,r). This indicates that the decreased staining intensity of TGN proteins in the Golgi of Vti1a/b DKO neurons is not caused by mislocalization to a different compartment in the soma.

Reduced staining intensity of TGN proteins could also be the result of epitope masking. To examine this possibility, TGN38-GFP was overexpressed in Vti1a/b DKO and control neurons and GFP fluorescence intensity in the Golgi was measured. The decrease in intensity of TGN38-GFP was similar to endogenous TGN38 (Sup. Fig. 2, Fig 1h). This rules out epitope masking as an explanation of reduced TGN38 staining intensity. Taken together, total levels of all analyzed Golgi proteins are decreased in the Golgi of Vti1a/b DKO neurons. While staining intensity of *cis*-/medial Golgi proteins is increased, the staining intensity of TGN proteins is decreased.

### Levels and localization of DCV markers are altered in Vti1a/b DKO neurons

We continued by analyzing proteins associated with DCVs. CgB and NPY-SEP are established endogenous and heterologous DCV markers, respectively (Arora et al., 2017; De Wit et al., 2009). As shown before, NPY-SEP intensity in the Golgi was not significantly different in the absence of Vti1a/b (Fig. 2m,n) (Emperador-Melero et al., 2018). In contrast, CgB staining intensity in the Golgi was increased (Fig. 2a,b).

**Fig. 2.**
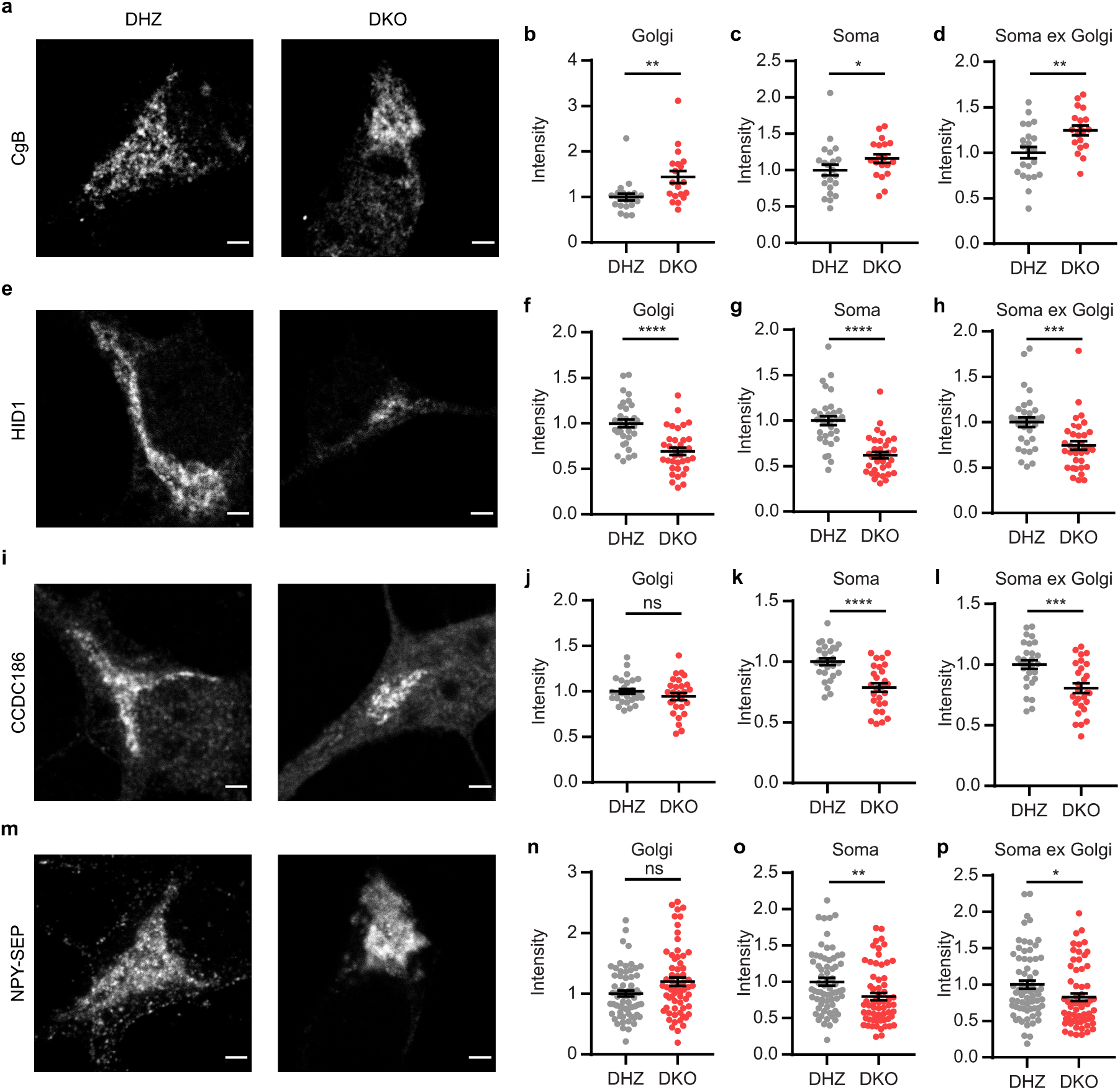
Levels and localization of DCV proteins are affected in Vti1a/b DKO neurons. **a** Representative examples of neurons immunostained for CgB. **b-d** CgB normalized staining intensity is increased in the Golgi (DHZ: 1 ± 0.0724, *n* = 22; DKO: 1.44 ± 0.132, *n* = 19; Mann Whitney test) (**b**), the soma (DHZ: 1 ± 0.075, *n* = 22; DKO: 1.16 ± 0.0605, *n* = 19; *t*-test) (**c**), and the soma excluding Golgi (DHZ: 1 ± 0.063, *n* = 22; DKO: 1.25 ± 0.0527, *n* = 19; *t*-test) (**d**). **e** Representative examples of neurons immunostained for HID1. **f-h** HID1 normalized staining intensity is decreased in the Golgi (DHZ: 1 ± 0.0434, *n* = 32; DKO: 0.694 ± 0.0398, *n* = 35; *t*-test) (**f**), soma (DHZ: 1 ± 0.0507, *n* = 32; DKO: 0.62 ± 0.0356, *n* = 35; Mann Whitney test) (**g**), and soma excluding Golgi (DHZ: 1 ± 0.0532, *n* = 32; DKO: 0.745 ± 0.0473, *n* = 35; Mann Whitney test) (**h**). **i** Representative examples of neurons immunostained for CCDC186. **j-l** CCDC186 normalized staining intensity is not significantly different in the Golgi (DHZ: 1 ± 0.0267, *n* = 28; DKO: 0.95 ± 0.0394, *n* = 27; *t*-test) (**j**), but decreased in the soma (DHZ: 1 ± 0.0394, *n* = 28; DKO: 0.786 ± 0.0353, *n* = 27; *t*-test) (**k**), and soma excluding Golgi (DHZ: 1 ± 0.0359, *n* = 28; DKO: 0.805 ± 0.0401, *n* = 27; *t*-test) (**l**). **m** Representative examples of neurons expressing NPY-SEP. **n-p** NPY-SEP normalized fluorescence intensity is not significantly different in the Golgi (DHZ: 1 ± 0.0484, *n* = 67; DKO: 1.2 ± 0.0701, *n* = 64; Mann Whitney test) (**n**), but decreased in the soma (DHZ: 1 ± 0.0509, *n* = 67; DKO: 0.797 ± 0.0491; Mann Whitney test) (**o**), and soma excluding Golgi (DHZ: 1 ± 0.0568, *n* = 67; DKO: 0.828 ± 0.0527, *n* = 64; Mann Whitney test) (**p**). Bars show mean ± SEM. *p < 0.05; **p < 0.01; ***p < 0.001; ****p < 0.0001. Scale bar is 2μm.

The increased CgB staining intensity suggests that endogenous DCV cargo accumulated in the Golgi. However, the total Golgi area was smaller in Vti1a/b DKO neurons (Fig. 1). Indeed, the integrated density (mean gray value multiplied by the total Golgi area) of both proteins was reduced (Sup. Fig. 1f,i). Hence, although CgB staining intensity was higher, the total amount of CgB staining (and NPY fluorescence) in the Golgi was in fact lower, arguing against cargo accumulation in the Golgi. In the soma (in general, and specifically the soma excluding the Golgi) NPY-SEP intensity was lower in Vti1a/b DKO neurons (Fig. 2o,p), while CgB staining intensity was higher (Fig. 2c,d). This shows that Golgi levels of DCV cargo are decreased in the absence of Vti1a/b, but that the effects on localization are different for CgB and NPY-SEP.

HID1 and CCDC186 play a role in the biogenesis/maturation of DCVs (Cattin-Ortolá et al., 2019; Hummer et al., 2017). Similar to TGN proteins, HID1 Golgi staining intensity was decreased in the Golgi, and in the whole soma (Fig. 2e-h). CCDC186 staining intensity was not different in the Golgi, while it was decreased in the whole soma (Fig. 2k,l). The integrated density of both proteins in the Golgi was reduced (Sup. Fig. 1g,h). Taken together, the levels and localization of proteins associated with DCVs are all altered by the absence of Vti1a/b, but the effects are different for different proteins.

### LAMP1 carrier staining is altered in Vti1a/b deficient neurons

Since the whole Golgi is disturbed by loss of Vti1a/b, we hypothesized that abnormal staining patterns are not restricted to DCVs and also exist for other types of vesicles, such as LAMP1 carriers. Indeed, compared to control neurons, Vti1a/b DKO neurons showed lower levels of LAMP1 staining intensity and the shape of these carriers was different (Fig. 3a,b). Control neurons contained characteristic oval shapes, while DKO neurons contained significantly fewer ovals (Fig. 3c,d). These results indicate that Vti1a/b are required for normal LAMP1 localization and levels.

**Fig. 3.**
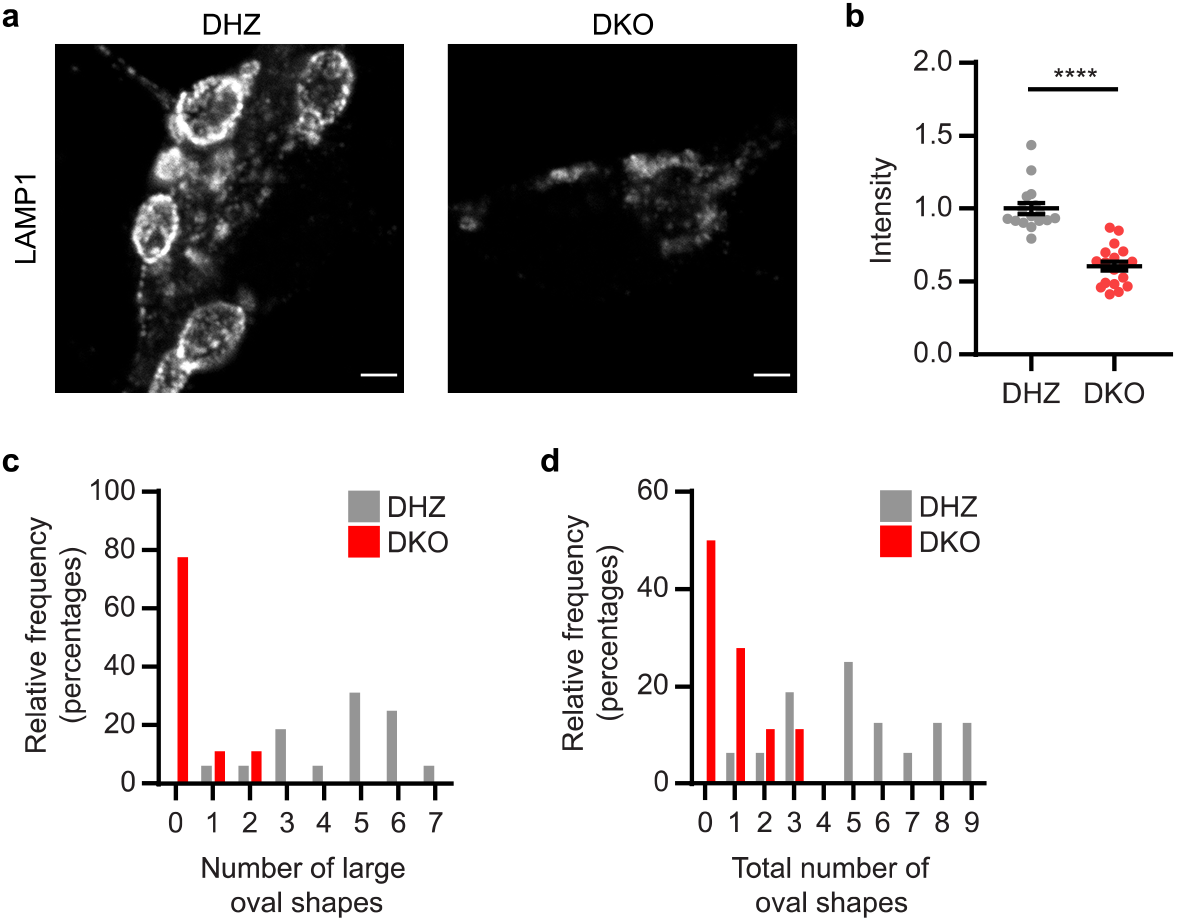
LAMP1 staining intensity and shape are abnormal in Vti1a/b deficient neurons. **a** Representative examples of neurons immunostained for LAMP1. **b** Normalized staining intensity of LAMP1 is decreased in Vti1a/b DKO neurons compared to DHZ neurons (DHZ: 1 ± 0.0397, *n* = 16; DKO: 0.606 ± 0.0326, *n* = 18; Mann Whitney test). **c** Number of large, clear oval LAMP1 shapes is decreased in Vti1a/b deficient neurons (DHZ: 4.5 ± 0.418, *n* = 16; DKO: 0.333 ± 0.162, *n* = 18; Mann Whitney test). **d** Total number of oval LAMP1 shapes (including less distinguished ovals) is decreased in Vti1a/b deficient neurons (DHZ: 5.31 ± 0.617, *n* = 16; DKO: 0.833 ± 0.246, *n* = 18; Mann Whitney test). Bars show mean ± SEM. *p < 0.05; **p < 0.01; ***p < 0.001; ****p < 0.0001. Scale bar is 2μm.

### Golgi abnormalities in Vti1a/b DKO neurons are not caused by reduced Golgi membrane availability or absence of the TGN compartment

Next, we investigated how loss of Vti1a/b causes disorganization of the Golgi apparatus, DCVs and LAMP1 carriers. We hypothesized that the absence of Vti1a/b leads to a smaller Golgi, which does not contain enough membrane to achieve normal Golgi function. In wildtype neurons, the Golgi grows significantly between DIV2 and DIV3 (Santos et al., 2017). Therefore, if membrane availability is the main cause of the Vti1a/b DKO phenotype, control neurons at DIV2 should show a similar phenotype to Vti1a/b DKO neurons. As expected, at DIV2, the Golgi area of control neurons was smaller than at DIV8 (Sup. Fig. 3a, Fig. 1b,e). The difference between control and DKO Golgi area at DIV2 was much smaller than the difference at DIV8. However, Golgi staining intensities of DKO neurons were affected similar to DIV8 neurons, when compared to controls (Sup. Fig. 3). This excludes membrane availability as a plausible cause for the Vti1a/b DKO phenotype.

Subsequently, we tested whether a TGN compartment exists in Vti1a/b DKO neurons. To test this, we overexpressed TGN38-GFP and stained for *cis*-Golgi marker GM130 and TGN protein TMEM87A. Line scans showed that the peak of intensity of TGN38-GFP was well separated from the peak of GM130 and close to the peak of TMEM87A (Fig. 4). No significant difference between control and Vti1a/b DKO neurons was observed. This suggests that TGN38-GFP can reach a TGN compartment in absence of Vti1a/b. Taken together, Golgi abnormalities in Vti1a/b DKO neurons are not caused by reduced Golgi membrane availability or the absence of a TGN compartment.

**Fig. 4.**
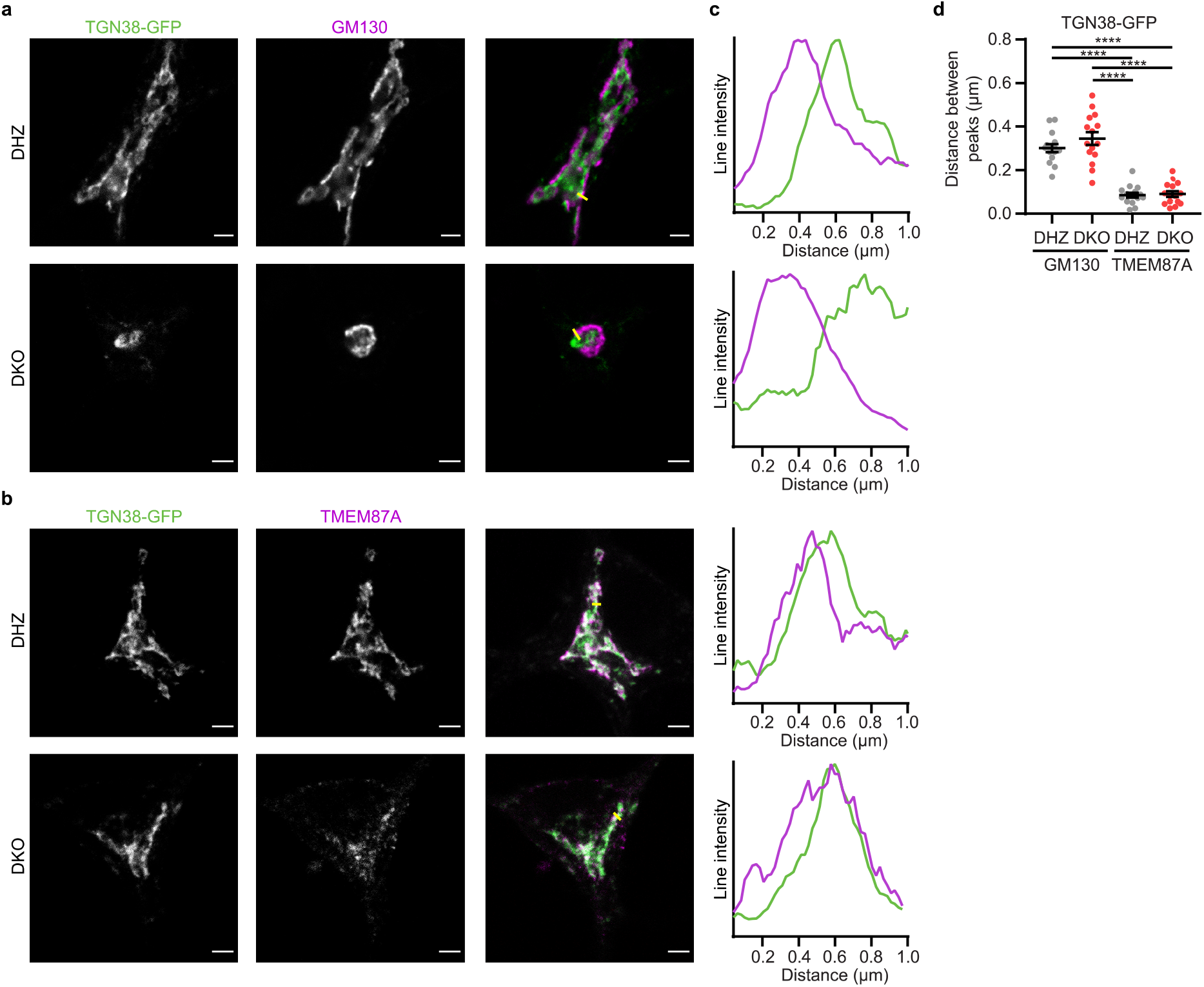
TGN38-GFP can reach the TGN compartment in Vti1a/b DKO neurons. **a** Representative examples of neurons overexpressing TGN38-GFP immunostained for GM130. (Brightness and contrast have been adjusted for visualization). **b** Representative examples of neurons overexpressing TGN38-GFP immunostained for TMEM87A. (Brightness and contrast have been adjusted for visualization). **c** Intensity profiles of TGN38-GFP and GM130/TMEM87A. Location of lines is represented as a yellow line in merged images. Intensities were normalized and smoothed before plotting. **d** The distance between intensity peaks of TGN38-GFP and GM130 (DHZ: 0.302 ± 0.0184, *n* = 15; DKO 0.346 ±0.0291, *n* = 15; ANOVA) is larger than the distance between peaks of TGN38-GFP and TMEM87A (DHZ: 0.0863 ± 0.0112, *n* = 15; DKO 0.09144 ± 0.013, *n* = 15; ANOVA) in both Vti DHZ and DKO neurons, measured by line scans through the Golgi. Bars show mean ± SEM. *p < 0.05; **p < 0.01; ***p < 0.001; ****p < 0.0001. Scale bar is 2μm.

### Vti1a/b DKO Golgi abnormalities are partially phenocopied by disturbing membrane homeostasis

Having ruled out reduced membrane availability as an explanation for the Vti1a/b DKO phenotype, we hypothesized that membrane homeostasis is disturbed in Vti1a/b DKO neurons. Previous research showed that addition of C6-Ceramide to HeLa cells and NRK cells results in distended Golgi cisternae, similar to what was observed in Vti1a/b DKO neurons (Emperador-Melero et al., 2018; Fukunaga et al., 2000; Hu et al., 2005). Further experiments showed that sphingosine, a metabolite of ceramide, causes these effects (Hu et al., 2005). We mimicked disturbed membrane homeostasis by addition of C6-Ceramide or L-Erythro-Sphingosine to wildtype neurons. C6-Ceramide can be metabolized, while L-Erythro-Sphingosine is a nonmetabolizable stereoisomer. After 4 hour incubation with C6-Ceramide or L-Erythro-Sphingosine, the Golgi was fragmented (Fig. 5a,b). Staining intensity for *cis*-Golgi marker GM130 was normal, but was decreased for both TGN proteins TGN38 and TMEM87A (Fig. 5). Since Golgi fragmentation was not observed in Vti1a/b DKO neurons, we conclude that disturbed membrane homeostasis cannot fully explain the Vti1a/b DKO phenotype, despite the fact that *cis*-Golgi and TGN proteins were differentially affected in both Vti1a/b DKO neurons and neurons treated with L-Erythro-Sphingosine or C6-Ceramide.

**Fig. 5.**
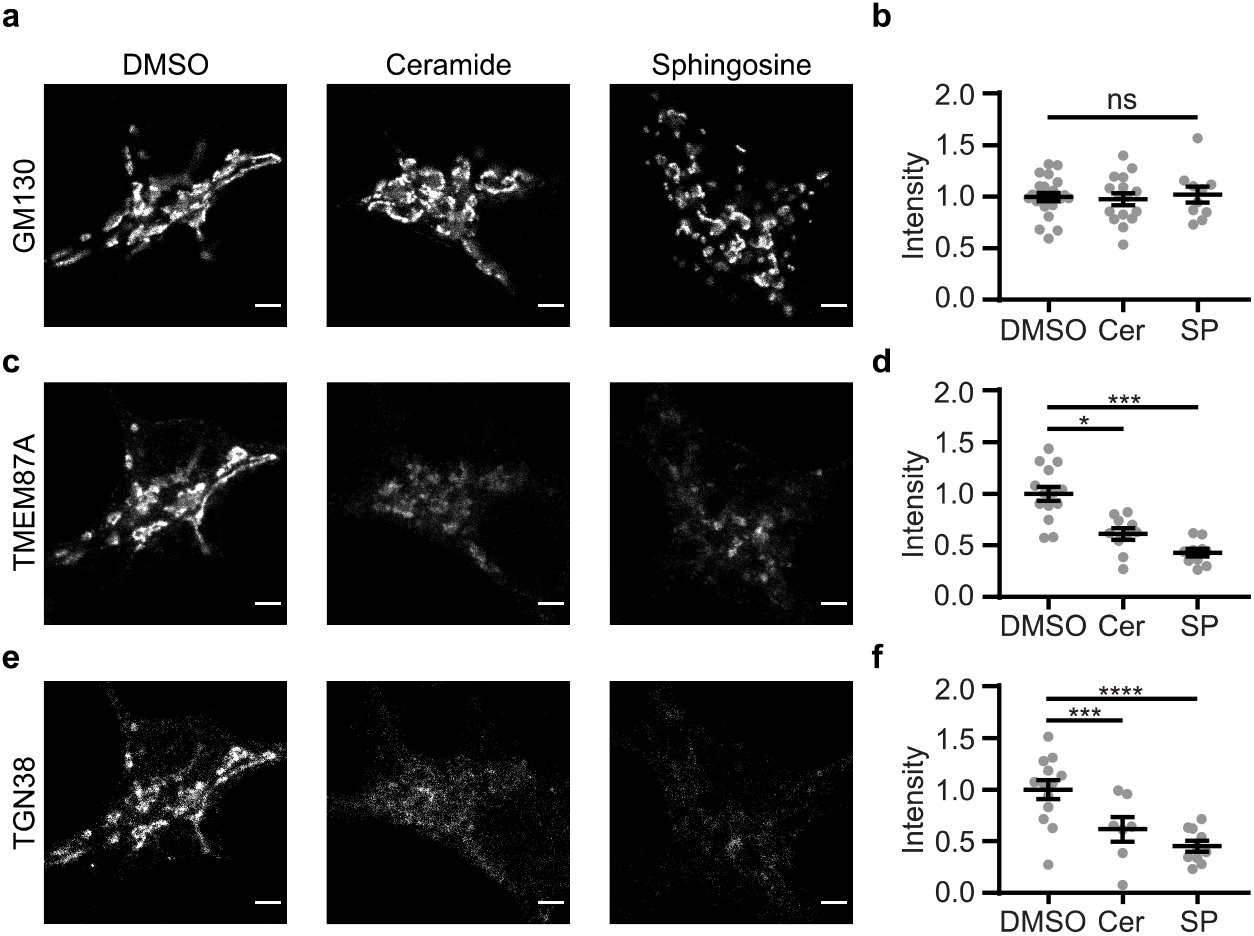
Addition of C6-Ceramide (Cer) or L-Erythro-Sphingosine (SP) to wildtype neurons does not affect staining intensity of *cis*-Golgi protein GM130, while TGN protein staining intensity is decreased. **a** Representative examples of neurons immunostained for GM130, after 4h treatment with 0.1%DMSO, 20 μM C6-Ceramide or 8 μM L-Erythro-Sphingosine. **b** GM130 normalized staining intensity is not affected by C6-Ceramide or L-Erythro-Sphingosine treatment (DMSO: 1 ± 0.0409, *n* = 22; Cer: 0.976 ± 0.0572, *n* = 16; SP: 1.02 ± 0.0775, *n* =10; ANOVA). **c** Representative examples of neurons immunostained for TMEM87A, after 4h treatment with 0.1% DMSO, 20 μM C6-Ceramide or 8 μM L-Erythro-Sphingosine. **d** TMEM87A normalized staining intensity is decreased after C6-Ceramide or L-Erythro-Sphingosine treatment (DMSO: 1 ± 0.0702, *n* = 14; Cer: 0.612 ± 0.0554, *n* = 10; SP: 0.43 ± 0.0374, *n* =10; ANOVA). **e** Representative examples of neurons immunostained for TGN38, after 4h treatment with 0.1% DMSO, 20 μM C6-Ceramide or 8 μM L-Erythro-Sphingosine. **f** TGN38 normalized staining intensity is decreased after C6-Ceramide or L-Erythro-Sphingosine treatment (DMSO: 1 ± 0.0906, *n* = 13; Cer: 0.615 ± 0.12, *n* = 7; SP: 0.453 ± 0.0522, *n* =10; ANOVA). Bars show mean ± SEM. *p < 0.05; **p < 0.01; ***p < 0.001; ****p < 0.0001. Scale bar is 3μm.

### Vti1a/b DKO Golgi abnormalities are not rescued by overexpression of sphingomyelin synthases or addition of myriocin

Similar to Vti1a/b DKO, deficiency of any of the components of the Conserved Oligomeric Golgi (COG) complex impacts retrograde trafficking, and morphology and function of the Golgi (Blackburn et al., 2019; Shestakova et al., 2006). Experiments in COG KD cells showed mislocalization of sphingomyelin synthase 1 (SMS1), the enzyme that converts ceramide into sphingomyelin (Spessott et al., 2010). It is conceivable that Vti1a/b are required for recycling of SMS1 to the Golgi. Two commercially available antibodies failed to produce reliable immunocytochemical stainings, precluding the analysis of endogenous SMS1 distribution in the absence of Vti1a/b. In addition to SMS1, which is mainly localized to the Golgi, sphingomyelin synthase 2 (SMS2) is found on the plasma membrane and in the Golgi (Huitema et al., 2004; Yeang et al., 2011). To visualize SMS1 and SMS2 in Vti1a/b DKO neurons, we tagged both with GFP. Similar to other TGN proteins, the intensity of SMS1-GFP and SMS2-GFP was lower in the Golgi of Vti1a/b DKO neurons as compared to Vti1a/b neurons, but no additional abnormalities were identified (Sup. Fig. 3a,b). If the Golgi phenotype in Vti1a/b deficient neurons was the result of mislocalization of SMS1 or SMS2, overexpression of these proteins could rescue the phenotype. However, while SMS1-GFP and SMS2-GFP were correctly targeted to the TGN, the Golgi area remained small and TGN protein intensity remained low (Sup. Fig 3). Thus, overexpression of SMS1-GFP or SMS2-GFP does not rescue the Vti1a/b DKO phenotype.

Deficiency of subunits of the GARP (Golgi-associated retrograde protein) complex also leads to retrograde trafficking defects and distended Golgi cisternae (Pérez-Victoria et al., 2008; Schmitt-John, 2015), resembling Golgi abnormalities in Vti1a/b DKO neurons. Myriocin is an inhibitor of sphingolipid synthesis and was shown to rescue GARP deficiency *in vitro* and *in vivo* (Fröhlich et al., 2015; Petit et al., 2020). We tested whether myriocin rescued Golgi abnormalities in Vti1a/b DKO neurons. Application of myriocin led to a trend towards an increased Golgi area at DIV7 (*p* = 0.159), and did not rescue the decreased TGN staining intensity in Vti1a/b DKO neurons (Fig. 1, Fig. 6). Hence myriocin does not rescue the Vti1a/b DKO phenotype.

**Fig. 6.**
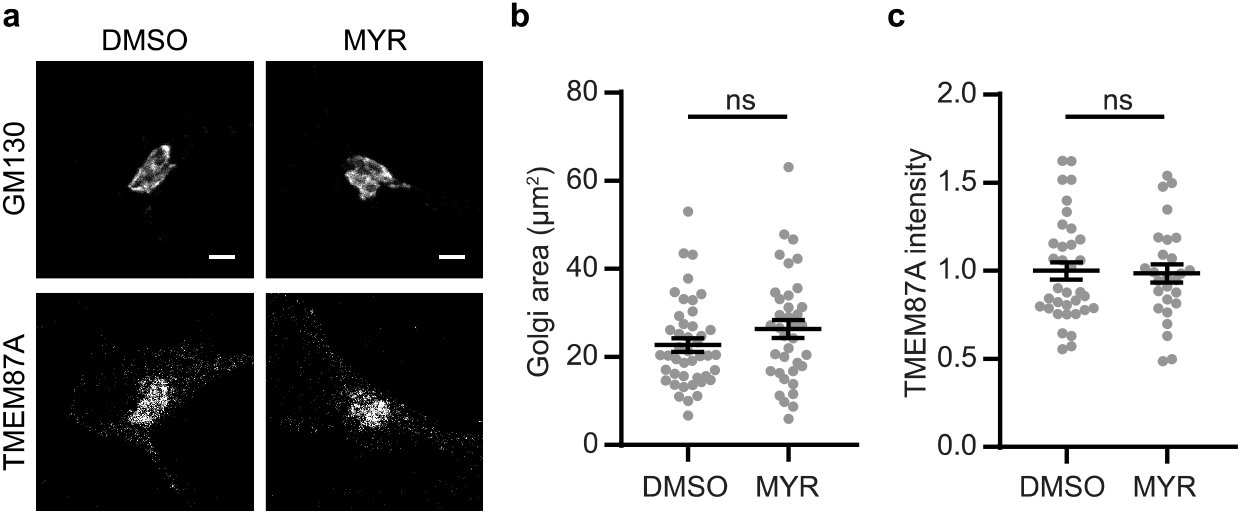
Addition of myriocin (MYR) to Vti1a/b DKO neurons does not rescue Golgi area or TGN staining intensity. **a** Representative examples of neurons immunostained for GM130 and TMEM87A, after 1 week treatment with 0.1% DMSO or 50 μM Myriocin. **b** Golgi area based on GM130 is not affected by myriocin treatment (DMSO: 22.7 ± 1.48, *n* =44; MYR: 26.4 ± 2.05, *n* =37; Mann Whitney test). **c** TMEM87A normalized staining intensity is not affected by myriocin treatment (DMSO: 1 ± 0.0503, *n* =35; MYR: 0.986 ± 0.0527, *n* =27; Mann Whitney test). Bars show mean ± SEM. *p < 0.05; **p < 0.01; ***p < 0.001; ****p < 0.0001. Scale bar is 3μm.

## Discussion

We investigated the role of retrograde trafficking in the organization of the Golgi apparatus and proteins involved in Golgi export. We showed that *cis*-/medial Golgi and TGN are affected differentially by the loss of Vti1a/b. This phenotype could not be explained by reduced Golgi membrane availability or absence of the TGN compartment, but was partially mimicked by addition of C6-Ceramide or L-Erythro-Sphingosine. The phenotype was not rescued by overexpression of sphingomyelin synthases or myriocin treatment.

### Condensed cis-/medial Golgi and diffuse TGN protein staining is unique for Vti1a/b DKO neurons

Loss of Vti1a/b results in reduced Golgi area in neurons (Emperador-Melero et al., 2018). For *cis*-/medial Golgi proteins, the integrated density (total area multiplied by the average staining intensity) was decreased, while at the same time the average intensity was increased. This suggests that the total protein level is lower, but the proteins that are present, are in a smaller area, causing a relative increase in intensity. TGN proteins showed both lower integrated density and average intensity in the Golgi. This indicates that total protein expression levels are decreased and this cannot be fully explained by a smaller Golgi area. Therefore, TGN proteins are more severely affected by loss of Vti1a/b than *cis*-/medial Golgi proteins.

This differential effect may be explained assuming that defective retrograde trafficking to the TGN affects TGN proteins more directly than *cis*-/medial Golgi proteins. If TGN proteins do not return to the TGN, they are expected to accumulate outside the Golgi. By measuring the staining intensity in the rest of the soma, we ruled out that proteins were accumulating there. It is possible that these proteins are degraded when they cannot return to the Golgi, resulting in decreased protein levels. Alternatively, epitope masking might explain decreased TGN marker levels. For example, a defect in protein maturation could result in the antibody not binding its epitope. Since overexpressed TGN38-GFP intensity was decreased to the same extent as endogenous TGN38, we excluded epitope masking as an explanation, at least for TGN38. In addition, the TGN38 antibody used in this study also detected overexpressed TGN38-GFP in Vti1a/b DKO neurons, indicating that the epitope is available in mutant neurons. Chia et al. performed an RNAi screen in HeLa cells for perturbations of Golgi morphology (Chia et al., 2012). They identified many genes of which depletion resulted in diffuse signals for *trans-Golgi* proteins, but in those cases *cis*-Golgi proteins were also affected to a similar extent. We are not aware of studies describing other gene depletions or treatments that result in reduced TGN staining but increased, condensed *cis*-/medial Golgi staining.

### Proteins associated with DCVs differentially depend on Vti1a/b

All proteins associated with DCVs showed decreased integrated density in the Golgi in absence of Vti1a/b, indicating that the levels of these proteins were reduced. As previously reported, the average intensity of NPY-SEP in the Golgi was not affected by loss of Vti1a/b, although a trend towards an increase was observed (Emperador-Melero et al., 2018). Golgi intensity of the endogenous DCV protein CgB was increased in the Golgi. Since protein levels were lower, this increase in average intensity was not caused by accumulation of CgB in the Golgi, but by reduction of Golgi size. While the intensity outside the Golgi apparatus was decreased for NPY-SEP, the intensity of CgB outside the Golgi increased. This could be explained by CgB being an endogenous and NPY-SEP a heterologous DCV marker, or it could be a difference between two DCV cargoes. It has been shown that CgB not only localizes to DCVs, but also to the nucleus in several cell types, including neuroendocrine cells, and plays a role in transcription (Yoo et al., 2002). Golgi stress in Vti1a/b DKO neurons could target CgB to the nucleus to induce changes in transcription. HID1 is thought to regulate DCV biogenesis in the TGN or DCV maturation by facilitating homotypic fusion (Du et al., 2016; Hummer et al., 2017). Like other TGN proteins, HID1 intensity inside and outside the Golgi was lower in Vti1a/b DKO neurons. This suggests that HID1 is indeed localized to the TGN. CCDC186 Golgi intensity was not affected, but the intensity outside the Golgi was decreased. Since CCDC186 is close to, but not exactly overlapping with a TGN marker (Cattin-Ortolá et al., 2019), it is logical that this protein is not affected in the same way as TGN proteins or DCV cargo. This is in line with our finding that loss of Vti1a/b differentially affected different compartments of the Golgi apparatus. Taken together, while the Golgi levels of proteins associated with DCVs were reduced in Vti1a/b deficient neurons, the impact on average staining intensities was different for each protein. This suggests that these proteins do not depend on Vti1a/b in the same way.

### Retrograde trafficking is required for proper Golgi morphology and function

The effects of Vti1a/b DKO are not specific for DCVs, since LAMP1 staining patterns were also disturbed in Vti1a/b deficient neurons. This is not surprising, considering that many different Golgi proteins are impacted by loss of Vti1a/b. Even though there is no significant difference in Golgi export of constitutive cargo in Vti1a/b DKO neurons at a late timepoint, the export is initially slower (Emperador-Melero et al., 2018). Thus, loss of Vti1a/b affects more cellular trafficking routes than regulated secretion. This suggests that defective retrograde trafficking impacts multiple secretory pathways. In line with this suggestion, disturbing retrograde trafficking by knocking out any of the four subunits of the GARP complex also disrupts anterograde trafficking and affects Golgi morphology (Hirata et al., 2015; Schmitt-John, 2015). In addition, knockout or knockdown of any of the components of the COG complex similarly affects Golgi morphology and function (Blackburn et al., 2019; Shestakova et al., 2006). Therefore, we conclude that disturbed retrograde trafficking affects Golgi morphology and function and thereby likely impacts many more Golgi-dependent processes than Golgi export of regulated cargo.

### Similar Golgi abnormalities upon disturbed membrane homeostasis and Vti1a/b deficiency appear not causally related

Disturbing membrane homeostasis by addition of ceramide or sphingosine produced similar Golgi abnormalities as observed in the absence of Vti1a/b. Disturbed membrane homeostasis can affect Golgi organization in different ways, for example by disrupting lipid rafts or by altering membrane permeability (Bieberich, 2018; Pascher, 1976; Tracey et al., 2018). Since Vti1a/b are required for retrograde trafficking, Vti1a/b deficiency may directly affect recycling of membrane components to the Golgi apparatus, but Golgi abnormalities may also be an indirect effect of impaired recycling of membrane-processing enzymes. We focused on sphingomyelin synthases because it was shown that retrograde trafficking defects cause mislocalization of SMS1 (Spessott et al., 2010). In addition, SMS1 and SMS2 are required for normal trafficking and secretion of insulin and vesicular stomatitis virus G protein tagged with GFP (VSVG-GFP) (Subathra et al., 2011). We did not observe abnormalities in localization of SMS1/2-GFP and SMS1/2 did not rescue Golgi abnormalities in Vti1a/b DKO neurons, nor did inhibition of *de novo* ceramide synthesis by myriocin application. This suggests that mislocalization of sphingomyelin synthases does not underly the defects in Vti1a/b deficient neurons. Since cells have many different ways to regulate ceramide levels (Hussain et al., 2012), we cannot exclude that the Vti1a/b DKO Golgi phenotype is the result of defects in other aspects of lipid metabolism, but clear leads, as in the case of SMS1/2 and myriocin, are lacking, to our knowledge. Altogether, we have shown that Vti1a/b are important for different aspects of TGN and *cis*-/medial Golgi organization.

## Methods

### Laboratory animals and primary cultures

All animals were bred and housed according to Institutional and Dutch governmental guidelines. Embryonic day (E) 18.5 Vti1a/b DKO, Vti1a/b DHZ and C57BL/6 mouse embryos were used for primary hippocampal culture. Vti1a/b DKO and DHZ mice have been described previously (Emperador-Melero et al., 2018; Kunwar et al., 2011). Mouse hippocampi were dissected in Hanks’ balanced salt solution (Sigma), supplemented with 10mM HEPES (Gibco) and were digested with 0.25% trypsin (Gibco) in Hanks’ + HEPES for 15 min at 37°C. Hippocampi were washed three times with Hanks’ + HEPES, once with DMEM complete (DMEM + Glutamax (Gibco), supplemented with 10% FCS (Gibco), 1% NEAA (Sigma) and 1% penicillin/streptomycin (Sigma)) and triturated with fire-polished Pasteur pipettes. Dissociated cells were spun down and resuspended in Neurobasal medium (Gibco) supplemented with 2% B-27 (Gibco), 1.8% HEPES, 0.25% Glutamax (Gibco) and 0.1% penicillin/streptomycin. Continental cultures were created by plating WT and Vti1a/b DHZ neurons at 25K/well and Vti1a/b DKO neurons at 50K/well. Neurons were seeded on pre-grown rat glia on 18mm glass coverslips in 12 well plates.

### Immunocytochemistry

Neurons were fixed at DIV2 or DIV7/DIV8 in freshly prepared 3.7% paraformaldehyde (EMS) for 10 min at room temperature and permeabilized with 0.1% Triton X-100 (Fisher Scientific) for 10 min. In case only goat secondary antibodies were used, neurons were blocked with 5% normal goat serum (Life Technologies) for 30 min. Incubation with primary and secondary antibodies was done at room temperature for 1 h. All solutions were in PBS (composition in mM: 137 NaCl, 2.7 KCl, 10 Na2HPO4, 1.8 KH2PO4; pH= 7.4). Coverslips were mounted in Mowiol (Sigma). In case donkey anti-sheep and goat antibodies were combined, secondary antibody incubation was split into two steps to prevent cross-reaction: donkey anti-sheep antibodies were incubated for 1h and after thorough washing and blocking with 5% normal goat serum, goat secondary antibodies were incubated for 1h.

Primary antibodies used for immunocytochemistry: GM130 (1:500, BD, 610822), giantin (1:200, BioLegend, 909701), TGN38 (1:100, BioRad, AHP499G), TMEM87A (1:100, Novus Biologicals, NBP1-90532), ARL1 (1:50, Santa Cruz, sc-393785), HID1 (1:100, Novus Biologicals, NBP2-02667), CCDC186 (1:200, Novus Biologicals, NBP1-90440), CgB (1:500, Synaptic Systems, 259 103), LAMP1 (1:500, Abcam, ab25245). Secondary antibodies were AlexaFluor-conjugated (1:500, Invitrogen) or STAR GREEN-, STAR580- or STAR635p-conjugated (1:100, Abberior).

For ceramide and sphingosine experiments, DIV7 neurons were treated with 20 μM C6-Ceramide (Cayman Chemical), 8 μM L-Erythro-Sphingosine (Cayman Chemical) or 0.1% DMSO (Sigma) for 4 hours. For myriocin experiments, 50 μM myriocin (Cayman Chemical) or 0.1% DMSO was added to neurons 4 hours after plating and neurons were fixed at DIV8.

Confocal imaging was performed on a Leica TCS SP8 STED 3x microscope (Leica Microsystems) with LAS X acquisition software, using a 100x oil objective (NA=1.4). A pulsed white light laser was used to excite the samples. The signal was detected using a gated hybrid detector (HyD) (Leica Microsystems) in photon-counting mode.

Images were analyzed in Fiji software (Schindelin et al., 2012). The BAR plugin was used to plot multichannel profiles in order to analyze the distance between peaks of Golgi proteins (Ferreira et al., 2015).

### Plasmids

NPY-SEP plasmid has been described before (van de Bospoort et al., 2012). TGN38-GFP, SMS1-GFP and SMS2-GFP were generated by amplifying TGN38, SMS1 and SMS2 from a mouse cDNA library and tagging them with eGFP at the C-terminus. The following primers were used for amplification of TGN38: gctcaccatgatcccaagctttaggttcaaacgttgg and cggtaccgcgggcccatgcggttccaggttgcg; SMS1: ggactcagatctcgaatgttgtctgccaggaccatgaaggaag and gaagcttgagctcgattatgtgtcgtttaccagccggctatatttaacttgcctactaaggt; SMS2: ggactcagatctcgaatggatatcatagagacagcaaaacttgaaggtcacttgg and gaagcttgagctcgatcaggtagacttctcattatcctccccgatcttctgg) These plasmids were cloned into lentiviral vectors under human Synapsin promoter and viral particles were delivered to neurons at DIV4 or DIV5 for NPY-SEP and DIV0 for SMS1-GFP and SMS2-GFP.

### Statistical analysis

Statistical analysis and graphing were performed using GraphPad Prism 7. Shapiro-Wilk was used to test distribution normality. When assumption of normality was met, parametric tests were used: *t*-test or ANOVA (with Tukey’s multiple comparisons test). Otherwise, non-parametric tests were used: Mann Whitney or Kruskall-Wallis (with Dunn’s multiple comparisons test).

## Supplementary figures

**Supplementary Figure 1:**
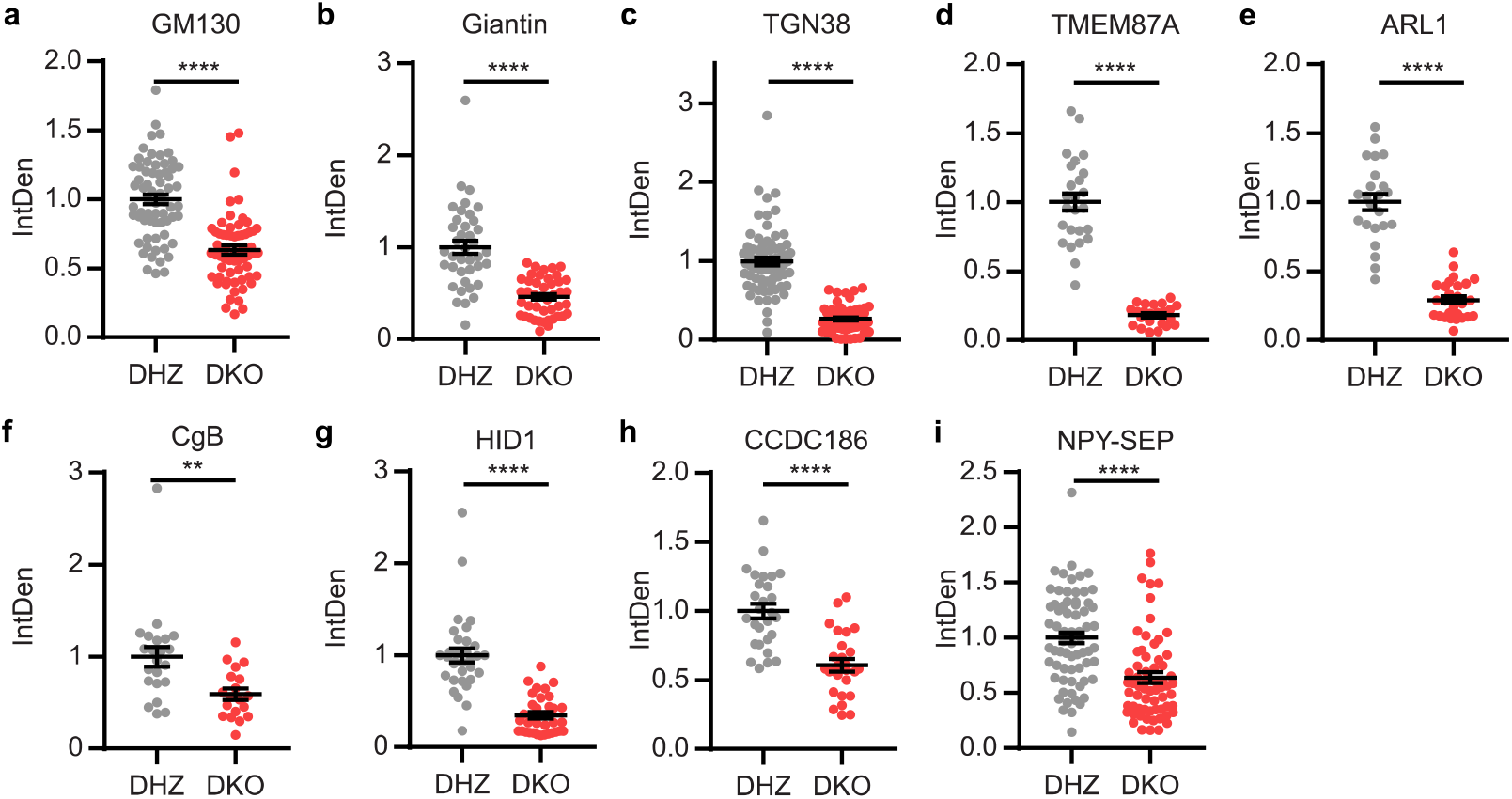
Golgi protein levels are lower in Vti1a/b DKO neurons. **a-i** Integrated density (IntDen; sum of all pixel values) of GM130 (DHZ: 1 ± 0.0356, *n* = 65; DKO: 0.631 ± 0.0331, *n* = 61; Mann Whitney test) (**a**), giantin (DHZ: 1 ± 0.0719, *n* = 39; 0.459 ± 0.0314, *n* = 44; Mann Whitney test) (**b**), TGN38 (DHZ: 1 ± 0.0482, *n* = 73; 0.267 ± 0.0187, *n* = 69; Mann Whitney test) (**c**), TMEM87A (DHZ: 1 ± 0.0628, *n* = 25; DKO: 0.181 ± 0.0153, *n* = 23; *t*-test) (**d**), ARL1 (DHZ: 1 ± 0.0609, *n* = 23; DKO: 0.291 ± 0.0265, *n* = 27; *t*-test) (**e**), CgB (DHZ: 1 ± 0.107, *n* = 22; DKO: 0.592 ± 0.0612, *n* = 19; Mann Whitney test) (**f**), HID1 (DHZ: 1 ± 0.0763, *n* = 32; DKO: 0.347 ± 0.0343, *n* = 35; Mann Whitney test) (**g**), CCDC186 (DHZ: 1 ± 0.0511, *n* = 28; DKO: 0.607 ± 0.0444, *n* = 27; *t*-test) (**h**) and NPY-SEP (DHZ: 1 ± 0.0502, *n* = 67; DKO: 0.64 ± 0.048, *n* = 64; Mann Whitney test) (**i**) in the Golgi of Vti1a/b DKO neurons, normalized to DHZ controls. Bars show mean ± SEM. *p < 0.05; **p < 0.01; ***p < 0.001; ****p < 0.0001.

**Supplementary Figure 2:**
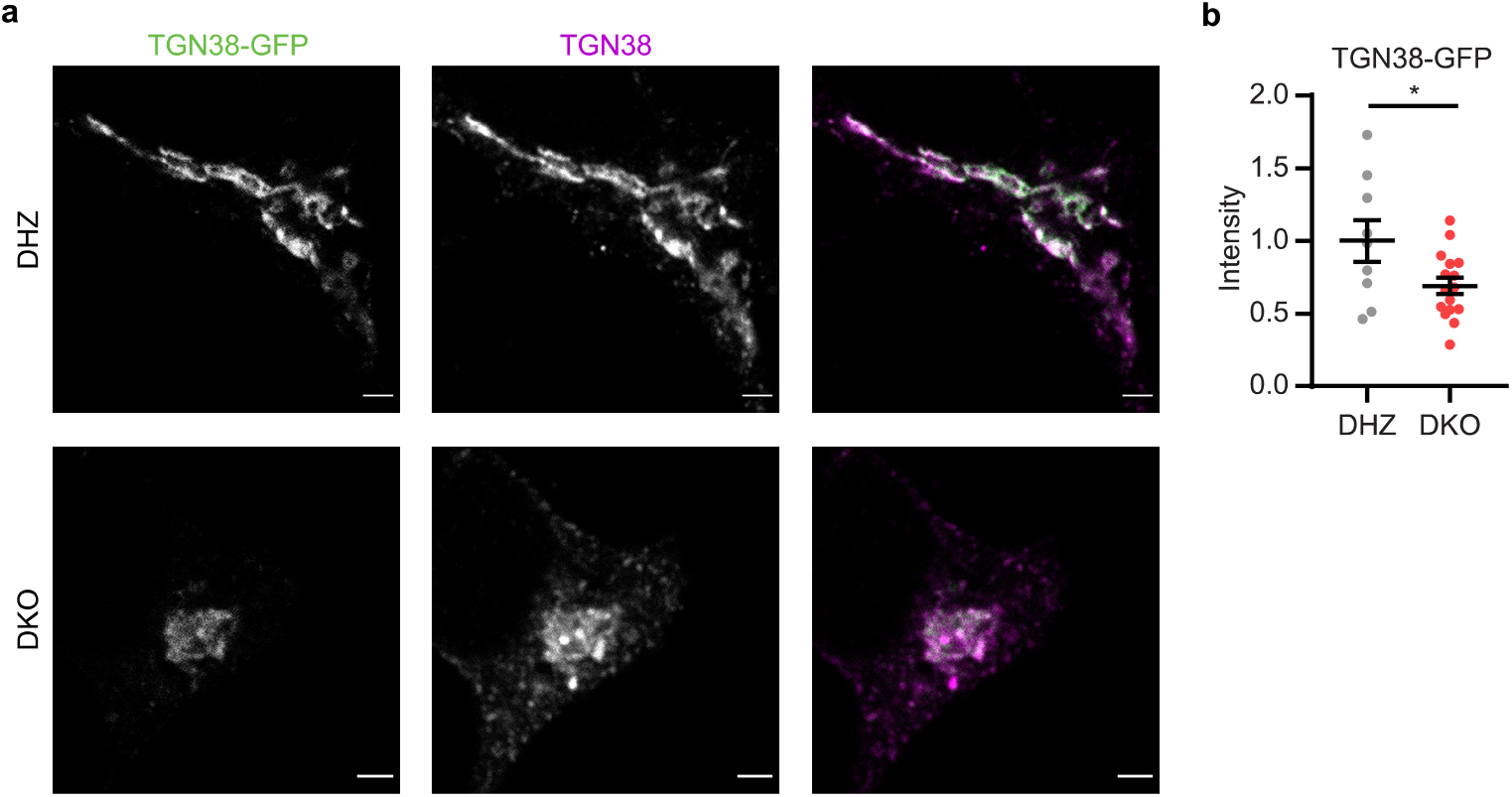
Decreased TGN staining intensity in Vti1a/b deficient neurons is not caused by epitope masking. **a** Representative examples of neurons overexpressing TGN38-GFP, immunostained for TGN38. **b** Normalized staining intensity of TGN38-GFP in the Golgi is decreased (DHZ: 1 ± 0.143, *n* = 9; DKO: 0.691 ± 0.057, *n* = 16; *t*-test). Bars show mean ± SEM. *p < 0.05; **p < 0.01; ***p < 0.001; ****p < 0.0001. Scale bar is 2μm.

**Supplementary Figure 3:**
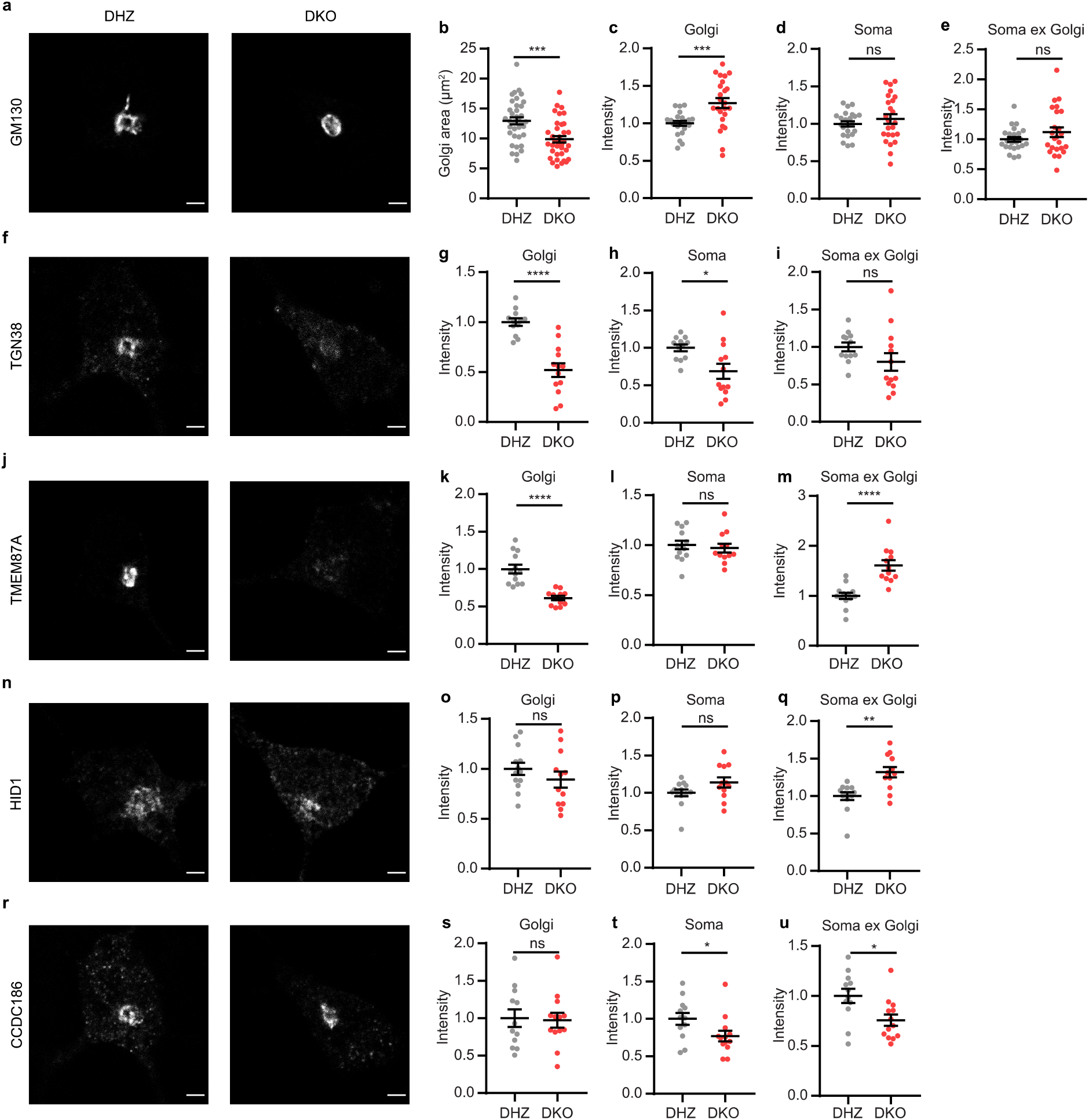
DIV2 Vti1a/b deficient neurons show similar phenotype as DIV8 neurons. **a** Representative examples of DIV2 neurons immunostained for GM130. **b** Decreased Golgi area in DIV2 Vti1a/b DKO neurons, compared to DHZ controls (DHZ: 13 ± 0.596, *n* = 36; DKO: 9.88 ± 0.526, *n* = 36; *t*-test). **c-e** GM130 normalized staining intensity is increased in the Golgi (DHZ: 1 ± 0.0313, *n* = 23; DKO: 1.27 ± 0.066, *n* = 24; *t*-test) (**c**) and not significantly different in the soma (DHZ: 1 ± 0.0347, *n* = 23; DKO: 1.07 ± 0.0638, *n* = 24; *t*-test) (**d**) and soma excluding Golgi (DHZ: 1 ± 0.0416, *n* = 23; DKO: 1.12 ± 0.0805, *n* = 24; *t*-test) (**e**). **f** Representative examples of DIV2 neurons immunostained for TGN38. **g-i** TGN38 normalized staining intensity is decreased in the Golgi (DHZ: 1 ± 0.0374, *n* = 12; DKO: 0.52 ± 0.0687, *n* = 13; *t*-test) (**g**) and soma (DHZ: 1 ± 0.0447, *n* = 12; DKO: 0.689 ± 0.0999, *n* = 13; *t*-test) (**h**), but not significantly different in the soma excluding Golgi (DHZ: 1 ± 0.0591, *n* = 12; DKO: 0.801 ± 0.116, *n* = 13; *t*-test) (**i**). **j** Representative examples of DIV2 neurons immunostained for TMEM87A. **k-m** TMEM87A normalized staining intensity is decreased in the Golgi (DHZ: 1 ± 0.0577, *n* = 13; DKO: 0.613 ± 0.0283, *n* = 12; *t*-test) (**k**), not different in the soma (DHZ: 1 ± 0.0439, *n* = 13; DKO: 0.969 ± 0.0445, *n* = 12; *t*-test) (**l**) and increased in the soma excluding Golgi (DHZ: 1 ± 0.0618, *n* = 13; DKO: 1.61 ± 0.106, *n* = 12; *t*-test) (**m**). **n** Representative examples of DIV2 neurons immunostained for HID1. **o-q** HID1 normalized staining intensity is not different in the Golgi (DHZ: 1 ± 0.0606, *n* = 13; DKO: 0.892 ± 0.081, *n* = 12; *t*-test) (**o**) and soma (DHZ: 1 ± 0.0472, *n* = 13; DKO: 1.14 ± 0.0672, *n* = 12; Mann Whitney test) (**p**) and increased in the soma excluding Golgi (DHZ: 1 ± 0.0511, *n* = 13; DKO: 1.32 ± 0.0696, *n* = 12; Mann Whitney test) (**q**). **r** Representative examples of DIV2 neurons immunostained for CCDC186. **s-u** CCDC186 normalized staining intensity is not different in the Golgi (DHZ: 1 ± 0.117, *n* = 12; DKO: 0.974 ± 0.0994, *n* = 13; *t*-test) (**s**) and decreased in the soma (DHZ: 1 ± 0.0806, *n* = 12; DKO: 0.772 ± 0.0713, *n* =13; *t*-test) (**t**) and soma excluding Golgi (DHZ: 1 ± 0.0718, *n* = 12; DKO: 0.758 ± 0.0567, *n* = 13; *t*-test) (**u**).Bars show mean ± SEM. *p < 0.05; **p < 0.01; ***p < 0.001; ****p < 0.0001. Scale bar is 2μm.

**Supplementary Figure 4:**
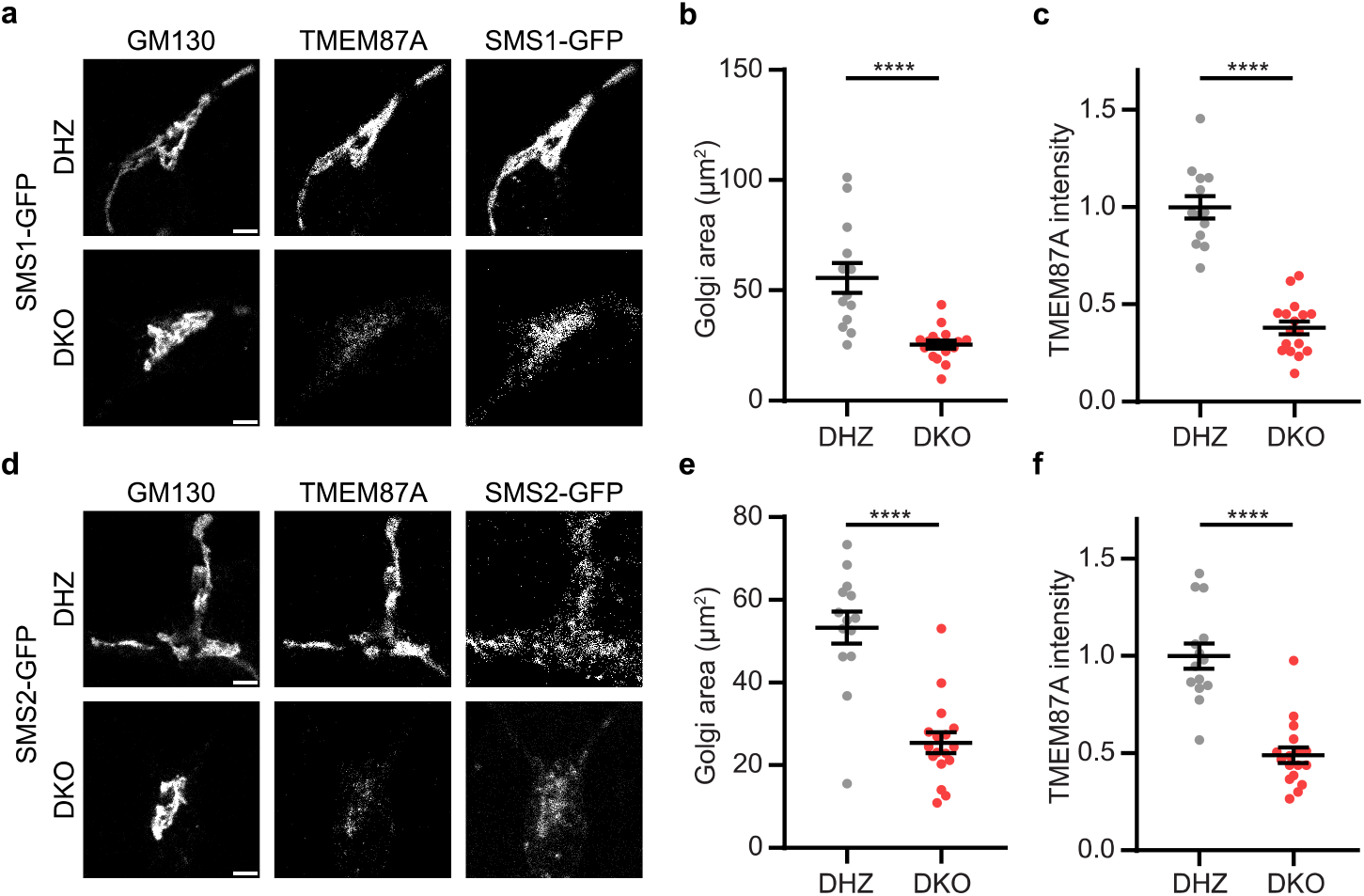
Overexpression of SMS1-GFP or SMS2-GFP does not rescue Golgi area or TGN staining intensity in Vti1a/b deficient neurons. **a** Representative examples of neurons overexpressing SMS1-GFP, immunostained for GM130 and TMEM87A. **b** Golgi area based on GM130 staining remains smaller in Vti1a/b DKO neurons overexpressing SMS1-GFP (DHZ: 55.8 ± 6.78, *n* = 13; DKO: 25.6 ± 1.81, *n* = 17; *t*-test). **c** Normalized staining intensity of TMEM87A in the Golgi remains lower in Vti1a/b DKO neurons overexpressing SMS1-GFP (DHZ: 1 ± 0.0567, *n* = 13; DKO: 0.38 ± 0.0329, *n* = 17; *t*-test). **d** Representative examples of neurons overexpressing SMS2-GFP, immunostained for GM130 and TMEM87A. **e** Golgi area based on GM130 staining remains smaller in Vti1a/b DKO neurons overexpressing SMS2-GFP (DHZ: 53.3 ± 3.85, *n* = 14; DKO: 25.5 ± 2.45, *n* = 17 *t*-test). **f** Normalized staining intensity of TMEM87A in the Golgi remains lower in Vti1a/b DKO neurons overexpressing SMS2-GFP (DHZ: 1 ± 0.0649, *n* = 14; DKO: 0.49 ± 0.0408, *n* = 17; *t*-test). Bars show mean ± SEM. *p < 0.05; **p < 0.01; ***p < 0.001; ****p < 0.0001. Scale bar is 3μm.

